# From one schema to another: How the prefrontal cortex responds to conflicting information

**DOI:** 10.1101/2025.08.08.669254

**Authors:** Petra Bíró, Silvy HP Collin

## Abstract

Over time, we develop event schemas or scripts that shape our expectations about what typically happens in certain contexts. However, even after forming a memory about a certain event, we are often exposed to related information about that same event at later points in time. This additional information sometimes causes one to have to re-evaluate the interpretation of the original event. Over a two-day fNIRS experiment, participants were exposed to events that were subsequently updated with schema-congruent or schema-incongruent additional details. These schema-incongruent additional details make those events more fitting to another schema than originally was the case, meaning that participants would need to dissociate that event from the original schema and re-integrate it with another schema. The fNIRS results showed stronger PFC activity for events updated with schema-congruent compared to schema-incongruent details. When specifically looking at those events that were updated with schema-incongruent details, our results suggest that dissociating an event from the original schema and re-integrating it with another schema was accompanied by an initial PFC decrease early in the trial followed by a PFC increase later in the trial. This was a distinctly different pattern compared to trials in which participants failed to re-integrate the event with another schema, which showed delayed PFC increase with lower amplitude and no initial PFC decrease. Our results refine our understanding of mechanisms of adaptive memory updating in the face of conflicting information.

## 1. Introduction

As we go through life, we encounter a wide range of experiences that help us develop event schemas—mental frameworks that guide which events we expect in particular situations. For instance, we might form a schema about typical events on holidays. A growing body of research has explored how these event schemas are represented in the brain (Bein and Niv, 2025; Beukers et al., 2024; Baldassano et al., 2018; Gilboa and Marlatte, 2017; Ghosh and Gilboa, 2014; van Kesteren et al., 2012), and how they aid memory encoding and help us remember certain events and everyday scenarios (Collin et al., 2025; Roy et al., 2025; Audrain and McAndrews, 2022; Masís-Obando et al., 2022; van Kesteren et al., 2020; Höltje et al., 2019; Greve et al., 2019; DuBrow et al., 2017; Brod et al., 2015; Schlichting and Preston, 2015; Van Kesteren et al., 2013). However, there is a possibility that the schemas in which we originally integrated certain events turn out to be false when new information comes to light. In that case, we would have to revise them, i.e., dissociate that certain event from its original event schema and reintegrate it with another schema. For example, imagine you read the following event description: People are gathered in a large building, where string lights hang overhead and the smell of cider and cookies fill the air. A jazz trio is playing softly in the corner as people mingle and take photos near a festively lit tree. Based on this information, you might interpret this as a holiday event. However, later you might learn that the large building is in fact a campus atrium, the jazz trio playing that day was composed of graduate students, and the festively lit tree was actually part of a visual attention study, tracking how festive stimuli influence spatial memory in a crowded environment. These additional details might make you re-evaluate this event from a holiday event to an educational event. Because of this, you might have to re-integrate your memory of this event with another event schema.

Earlier research examined how episodic memories are corrected or updated with new information (Bein et al., 2023; Kiley and Parks, 2022; Carpenter et al., 2021; Sinclair and Barense, 2018; St. Jacques and Schacter, 2013; Hupbach et al., 2008; Forcato et al., 2007). These and other studies generally conclude that updates of memories mostly take place at moments of re-consolidation, i.e., right after a consolidated memory is reactivated and susceptible to change (Richter et al., 2019; Schiller and Phelps, 2011; Schiller et al., 2010). However, these memory updating studies concern primarily about how and when to update episodic information *itself*. Less is known about the neural mechanisms behind updating the attachment of episodic memories from one to another schema context due to new information coming to light.

How does the brain support our ability to determine schema-congruency of events in the first place? Previous findings suggest that the prefrontal cortex (PFC) is strongly involved in terms of our ability to determine schema-congruency of information, with most studies showing medial PFC involvement for schema-congruent information (Bein and Niv, 2025; Guo and Yang, 2023; Masís-Obando et al., 2022; Gilboa and Marlatte, 2017; van Kesteren et al., 2012) and dorsolateral PFC involvement when information is somehow incongruent (Ohashi et al., 2018; Brod et al., 2015; Plaza et al., 2008). Building on earlier research, we aim to investigate the role of the PFC in situations where information is not inherently false, but was initially integrated into a schema based on incorrect assumptions or information. Then, when this new information reveals that the information is integrated into the wrong schema, the previously integrated content must be dissociated from its original schema context and (re)integrated into a more appropriate one.

Due to our focus on the PFC in this study, we used low-density functional near-infrared spectroscopy (fNIRS) with optodes situated to continuously measure the PFC. Many studies on the neural basis of schema-mediated memory are focused on functional magnetic resonance imaging (fMRI) instead, which has the advantage of high spatial resolution and full brain coverage. However, it also some downsides in some types of cognition research, like its high sensitivity to motion and the fact that you have to place the participant in a loud and strongly confined location. Low-density fNIRS is more portable and less sensitive to motion. Therefore, fNIRS might be ideally suited for studies in young children, clinical populations or studies that use naturalistic tasks. However, given the strong focus on fMRI in schema-mediated memory research, it is important to first replicate some earlier findings from fMRI research but now using fNIRS (Pinti et al., 2020), before moving on to our main goal of investigating the role of the PFC in re-integrating events from one to another schema context.

Prior research has already replicated some previous fMRI findings using fNIRS. For example by showing significant dorsolateral PFC activity during encoding of new episodic memories (novel faces in this case) (Jahani et al., 2017), showing greater PFC activity during more demanding working memory tasks (Boere et al., 2024; Yang et al., 2024b; Llana et al., 2022; Lucas et al., 2020), showing greater PFC activity during a prospective memory task (Koo et al., 2022), and showing medial PFC activity during intentional forgetting compared to intentional remembering in a directed forgetting paradigm (Jing et al., 2022). However, there is a lack of fNIRS research showing more (medial) PFC involvement when confronted with schema-congruent information which is a well established finding from fMRI work (Bein and Niv, 2025; Gilboa and Marlatte, 2017; van Kesteren et al., 2012). For this reason, before moving on the our main research question, we want to answer the following initial question: *Is (m)PFC activity as measured with fNIRS higher for schema-congruent compared to schema-incongruent information?* Then, we will focus on our main research question for this study: *How does the PFC contribute to dissociating an event from one schema and re-integrating it into another when new information prompts a shift in schema affiliation?* To investigate these open questions, this study will measure PFC activity using fNIRS during encoding of schema-related events that are, on a subsequent day, updated with additional details that are either consistent or not consistent with their original schema. We predicted higher activity in channels mapping onto medial PFC for events that are only updated with schema-congruent details, compared to event that were updated with schema-incongruent details. To furthermore investigate how we dissociate events with their original schema and subsequently re-integrate events with another schema when new information comes to light, we will split the events that were updated with schema-incongruent information into those for which a participant subsequently successfully changed its schema-affiliation (referred to as *schema-updating events*) compared to those for which a participant’s memory remains based on the old schema-affiliation (referred to as *schema-violating events*). By comparing across schema-updating, schema-violating, and control events that are updated with only schema-consistent information (referred to as *schema-consistent events*), we can shed more light onto the exact contribution of the PFC in encoding schema-incongruent information, by examining how activity across PFC sub-regions differs when participants successfully accommodate new but schema-incongruent information versus when they fail to do so.

## 2. Methods

### 2.1. Participants

38 participants took part in the experiment, comprising 24 women and 15 men. Participants ranged in age from 18 to 33 years (M = 21.49, SD = 3.42). Recruitment was conducted via convenience sampling using Tilburg University’s Sona participant pool, and data acquisition was done at the DAF Technology lab (Tilburg University). The participants received credits through Sona for their participation. Participants gave written informed consent before starting the experiment. Two participants were excluded due to technical issues during data acquisition. The experiment was approved by the Research Ethics and Data Management Committee of TSHD, Tilburg University.

### 2.2. Stimuli

For the study, we created 32 descriptions of fictional activities. Each activity was part of either a holiday or a education schema. For each description, participants were made aware of whether this description was part of the holiday or the education schema by showing them a map of a fictional holiday Island (Figure 1A) or of a fictional college campus (Figure 1B). Each description was specifically linked to 1 of 9 possible locations at that map by circling the correct location for that activity description. Participants read the description alongside seeing the map with the circled location, with the description being placed below the map on the screen, and a general title for the activity description above the map. All used descriptions were created using chatGPT followed by manual changes to finetune them.

**Figure 1:**
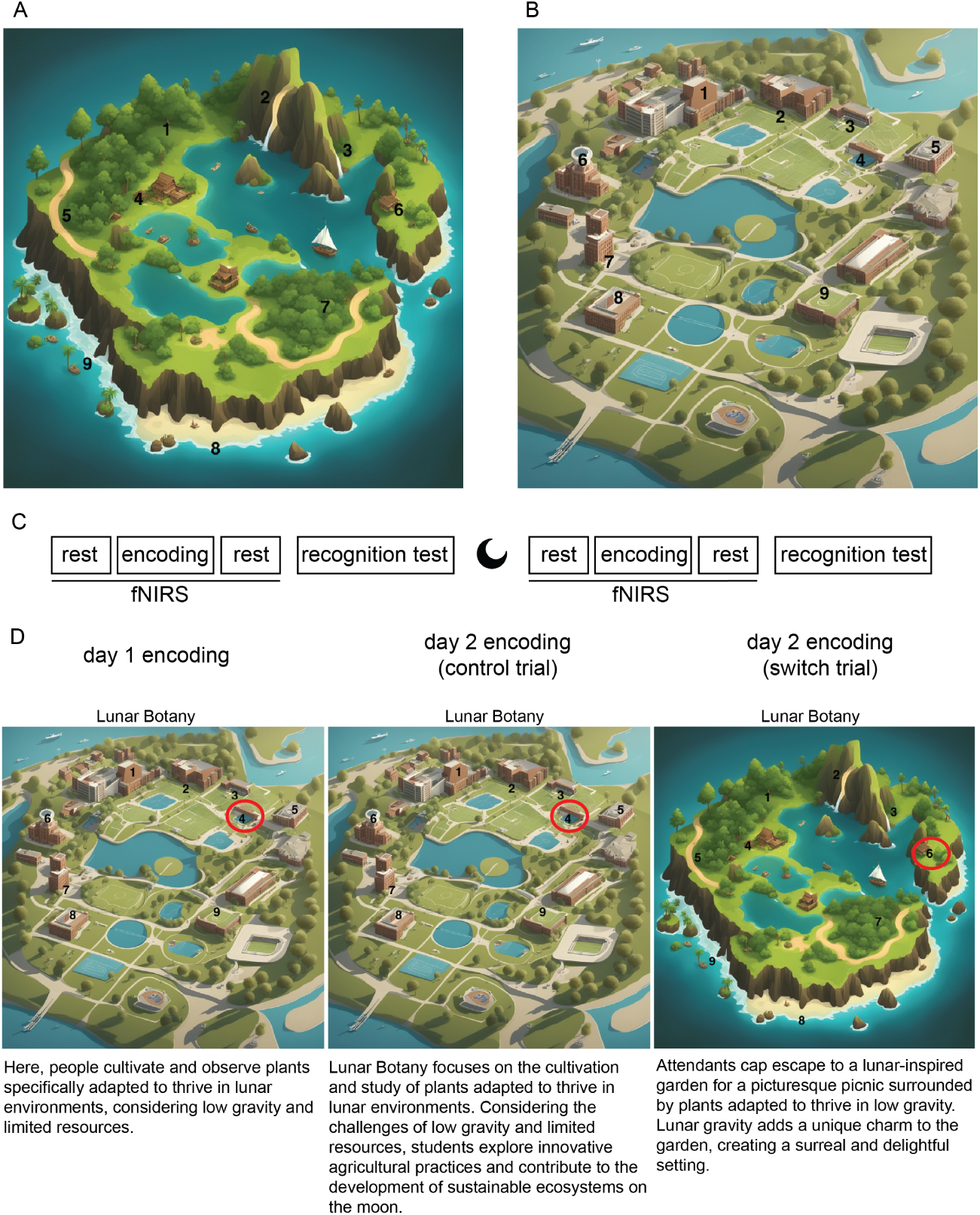
Overview of stimulus material and experimental design. **A:** The map used for the holiday schema. **B:** The map used for the education schema. **C:** Overview of the entire experiment. **D:** From left to right: Example stimulus of day 1 encoding, example stimulus of day 2 encoding when that activity description is a control trial, example stimulus of day 2 encoding when that activity description is a switch trial.

**Figure 2:**
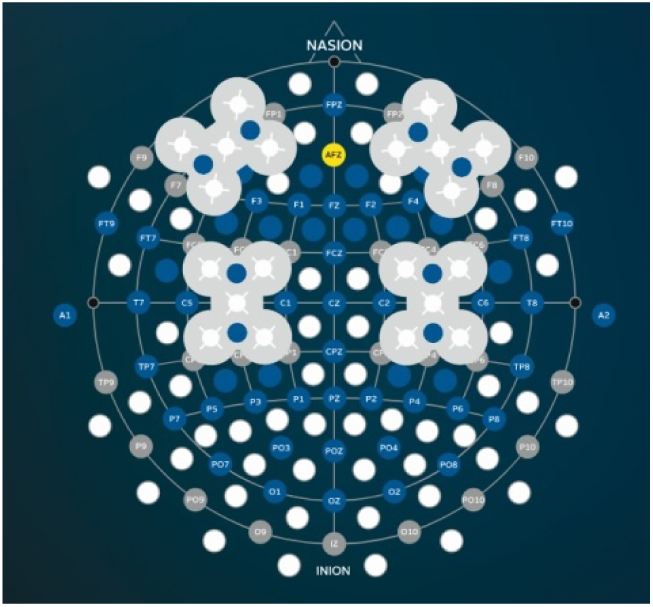
fNIRS transmitter placement. © g.tec medical engineering GmbH, 2025. Used with permission of the rights holder. The eight fNIRS transmitters were approximately located at AFF4h, F6, AF8, AFp4, AFF3h, F5, AF7, and AFp3 positions, while two receivers were placed at AF7 and AF8 (Schreiner et al., 2021).

### 2.3. Experimental procedure

#### 2.3.1. Day 1

Participants were seated in front of a computer monitor wearing a g.tec fNIRS headcap (g.tec, n.d.), and instructed to remain as still as possible and follow the on-screen instructions. It started with a 3 minute rest block in which they were instructed to not think of anything in particular, which was followed by the encoding task. During the encoding task, they received 32 activity descriptions, one at the time, in a random order. Each activity description was associated with a map and a particular location on that map, by showing the activity description below that map, and by circling in red the particular location at which that activity happened on the map (Figure 1D). This was repeated 6 times. Each stimulus was presented for 9 seconds, followed by a 1-second fixation. Participants viewed 16 holiday-related and 16 educational activities, presented in random order. After the entire encoding task finished, they again had a 3 minute rest block. The last task was the recognition test which measured if and how well they remembered these stimuli with three questions for each of the 32 activity descriptions: whether the activity was old (i.e., seen before) or new (i.e., not seen before), which schema/map it belonged to (holiday or education), and which location did it happen (from 9 possible locations on that map). Participants were also asked to rate their certainty for each of these 3 questions on a scale from 1 to 4. The memory test included old, lure and completely new descriptions; lure activities differed from the original ones by only a single word (e.g., Original: Here, people cultivate and observe plants specifically adapted to thrive in lunar environments, considering low gravity and limited resources. Lure: Here, people cultivate and observe trees specifically adapted to thrive in lunar environments, considering low gravity and limited resources.), while neutral activities were completely unrelated (e.g., In the community garden, residents cultivate not only plants but also connections, exchanging gardening tips, fostering a sense of communal growth and shared green-thumb triumphs). The descriptions were randomized and the task was self-paced. Both 3 minute rest-blocks and the encoding task included wearing the g.tec fNIRS headcap, the headcap was taken off before the recognition test. The encoding task was presented on a computer screen using the E-Prime 3.0 presentation software (Psychology Software Tools, Inc., 2016). The recognition test was administered using Qualtrics (Qualtrics, 2020).

#### 2.3.2. Day 2

Day 2 was identical to day 1, however, now the activity descriptions from day 1 were longer and richer, meaning participants had to update their memory for the day 1 activity descriptions with more details (for example, day 1 description: Here, people cultivate and observe plants specifically adapted to thrive in lunar environments, considering low gravity and limited resources. Day 2 description: Lunar Botany focuses on the cultivation and study of plants adapted to thrive in lunar environments. Considering the challenges of low gravity and limited resources, students explore innovative agricultural practices and contribute to the development of sustainable ecosystems on the moon. See Figure 1D). For both schemas/maps, 8 descriptions were control trials (i.e., schema congruent, remained an activity location at the same map/schema) and 8 were switch trials (i.e., schema incongruent, activity switched from one map/schema to the other). On day 2, the stimuli were repeated four times. Each stimulus was shown for 14 seconds, followed by a 1-second fixation. The encoding task was presented on a computer screen using the E-Prime 3.0 presentation software (Psychology Software Tools, Inc.,

2016). The recognition test was administered using Qualtrics (Qualtrics, 2020).

### 2.4. Equipment

#### 2.4.1. fNIRS data collection

The g.Nautilus fNIRS system was used, which supports wireless recording of fNIRS signals (g.tec, n.d.). Eight low-power optodes were positioned over the frontal cortex, and data were sampled at a frequency of 250 Hz (see Figure 2). The device utilizes LED technology, emitting light at standard wavelengths of 760 nm and 850 nm (g.tec, n.d.). Data acquisition was synchronized via Bluetooth between the g.tec system and Simulink/Matlab software.

### 2.5. Data pre-processing

#### 2.5.1. fNIRS data

fNIRS data were collected on both days of the experiment, resulting in two.mat files per participant that were read into Python using H5py (3.11.0) (h5py Developers, 2024). Each .mat file contains time series data from 16 channels, recording oxy-hemoglobin (HbO) and deoxy-hemoglobin (HbR) levels during the encoding phase, as well as a channel that indicated when the stimuli were presented. fNIRS data preprocessing was carried out using MNE (1.9.0) (Gramfort et al., 2013), along with numpy (1.26.4) (Harris et al., 2020) and pandas (2.2.2) (McKinney, 2010). For visualization, matplotlib (3.9.2) was used (Hunter, 2007). The .mat files created by g.tec during data acquisition contained concentration level data, there was no need to convert raw optical density data to concentration levels.

Following previous studies that applied fNIRS pre-processing steps and fNIRS MNE package tutorials (Gramfort et al., 2024; Jahani et al., 2017; Schreiner et al., 2021; Tak and Ye, 2014), the following fNIRS data processing pipeline was used. An MNE info object was created to include channel data (16 channels alternating between HbO and HbR data), stimulus information, and a frequency of 250 Hz. Subsequently, an MNE raw object was created for each participant. The raw data were bandpass filtered between 0.01 Hz and 0.1 Hz using a zero-phase finite impulse response (FIR) filter (low transition bandwidth: 0.02 Hz; high transition bandwidth: 0.005 Hz) to remove slow signal drifts and high-frequency physiological noise such as cardiac and muscle activity (Gramfort et al., 2024). MNE events and epoch objects were then created, with epochs for day 1 ranging from 0 to 8 seconds, and for day 2 from 0 to 13 seconds. The epochs were downsampled to 10 Hz (Montero-Hernandez et al., 2018), resulting in 10 samples per second. We performed a global baseline correction on the fNIRS data by subtracting the average of the rest-block prior to the task (using the last 2.5 minutes of this 3 minute rest block) from the epochs. These data were imported into R for statistical analysis. Appendix Figure A.3 showed the mean difference for HbO and HbR levels across all included trials on day 2.

#### 2.5.2. E-prime data

E-prime data, which contained the random sequence of activity descriptions for each participant, was exported using E-Data Aid as an Excel file and then converted into a .csv file using Microsoft Excel. This file was subsequently imported into Python using the Pandas package. The sequence data was then incorporated as epoch metadata with the MNE package.

### 2.6. Statistics

Statistical analyses were conducted using R software (version 4.2.2) (R Core Team, 2024), within the RStudio IDE (version 2023.09.0+463). For data pre-processing, visualization, and descriptive statistics, the following packages were used: psych (2.3.6) (Revelle, 2024), dplyr (1.1.4), tidyr (1.3.1), readr (2.1.4), stringr (1.5.0), and ggplot2 (3.4.4) (Wickham, 2019). For inferential statistics and modeling, the lmerTest (3.1.3) (Kuznetsova et al., 2017) and emmeans (1.11.0) (Lenth, 2024) packages were used. The final statistical outputs and formatting were completed using Quarto (version 1.4.553) (Posit, 2024), along with the tinytex (0.57) (Xie, 2023) and xtable (1.8.4) (Baird, 2021) packages for LaTeX integration.

#### 2.6.1. Statistics for comparing control to switch trials

Mixed-effects linear regression was applied (Pinheiro and Bates, 2000) to assess whether the impact of trial type (control vs. switch) on HbO concentration levels during encoding is moderated by the day of data collection (day 1 or day 2). Epochs from both days were annotated with their respective trial types (control or switch). Trials from day 2 were excluded if participants failed to correctly answer to the old/new question from the recognition test, meaning we only took stimuli into account that participants actually remembered. A random intercept was included for each participant to account for repeated measurements. The dependent variable, *value*, represents the continuous HbO signal, while the independent variables included *day* ( day 1 or day 2) and *trial type*, which categorizes the trials as schema congruent (i.e., control) or schema violating (i.e., switch), see equation 1). This was again done on the aggregated data (i.e., from all channels combined).

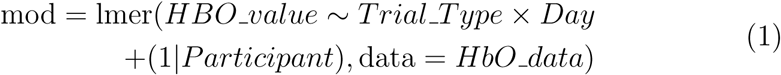

The same mixed-effects models were run for three different PFC region of interests (ROIs), to investigate whether the influence of schema congruency on HbO levels varies across different PFC sub-regions. The fNIRS channels were grouped into specific ROIs; parts of the medial prefrontal cortex (mPFC), parts of the inferior frontal gyrus (IFG), and parts of the dorsolateral prefrontal cortex (dlPFC) based on the labeling seen in Table 1 (Pfurtscheller and Da Silva, 1999; Schreiner et al., 2021).

**Table 1:** Optode, Electrode location, and Likely Region.

| Optode | Electrode Location | Likely Region |
| --- | --- | --- |
| 1 | AFp4 | mPFC / vmPFC |
| 2 | AF8 | Inferior Frontal Gyrus |
| 3 | F6 | Dorsolateral PFC |
| 4 | AFF4h | Dorsolateral PFC |
| 5 | AFp3 | mPFC / vmPFC |
| 6 | AF7 | Inferior Frontal Gyrus |
| 7 | F5 | Dorsolateral PFC |
| 8 | AFF3h | Dorsolateral PFC |

#### 2.6.2. Statistics for comparing schema-consistent to schema-updating and schema-violating trials

Mixed effects linear regression (Pinheiro and Bates, 2000) was used to investigate the difference in fNIRS signal for schema-updating compared to schema-violating trials on day 2. This model includes only successfully retrieved trials, which are determined as those, where the schema was remembered on day 1 (i.e., correct answer for the old/new question and the question about holiday or education) and all 3 questions were answered correctly on day 2. These trials were split into 3 congruency categories:

- Schema consistent trials, which only contains control trial data
- Schema updating trials, which contains switch trials where the performance on the recognition test on day 2 was correct according to the new, day 2 information
- Schema violating, which contains switch trials where the performance on the recognition test on day 2 was correct according to old, day 1 information.

The dependent variable in these models is *value*, which represents the HbO brain data, a continuous variable. The independent variable is *congruency*, a categorical variable with three levels: schema consistent, schema updating, and schema violating (see equation 2).

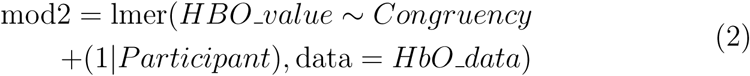

Given the delayed peak of the BOLD-related fNIRS response, all these mixed-effects models (from both sections 2.6.1 and 2.6.2) were limited to the time window beginning at 4 seconds post-stimulus onset to better capture event-related neural activity. For this reason, we placed a shaded grey area (0–4 s post-stimulus) on each lineplot that indicates the time window excluded from analysis to account for the latency of the hemodynamic response function (HRF) that underlies fNIRS measurements.

#### 2.6.3. Outliers

We excluded 2 participants from all analysis due to excessive motion during the experiment. Additionally, before statistical modeling, we determined participants who were outliers based on their HbO activity increase from baseline being more than 2 standard deviations away from the group mean and excluded those from analysis. For the analysis described in section 2.6.1, this excluded 2 additional participants. For the analysis described in section 2.6.2, this excluded one additional participant.

## 3. Results

For the old/new recognition question, participants performed well above 50%-chance level on day 1 (mean = 0.77, SD = 0.42) and day 2 (mean = 0.72, SD = 0.45). Table 2 overviews the behavioral results of the recognition tests, split for control and switch trials. For switch trials, these results suggest that people more often remembered the original schema and location of the story than the new one from day 2. This suggests that participants had a hard time switching to the other schema for the switch trials, and instead of switching, they often reported the original schema instead.

**Table 2:** Mean recognition test scores on day 2, split into trial type (control vs switch) and type of answers. When accDay1 is part of the description in the “Day” column, participants were correct on day 2 according to day 1 information. The question about which map has a 50% chance-level and the question about which location has a 11% chance-level.

|  | Trial.Type | Question | Day | Mean | SD |
| --- | --- | --- | --- | --- | --- |
| 1 | control | which map (holiday or education) | Day2 | 0.76 | 0.43 |
| 2 | control | which map (holiday or education) | Day2_accDay1 | 0.76 | 0.43 |
| 3 | control | which location (of 9 options) | Day2 | 0.63 | 0.48 |
| 4 | control | which location (of 9 options) | Day2_accDay1 | 0.63 | 0.48 |
| 5 | switch | which map (holiday or education) | Day2 | 0.21 | 0.41 |
| 6 | switch | which map (holiday or education) | Day2_accDay1 | 0.45 | 0.50 |
| 7 | switch | which location (of 9 options) | Day2 | 0.16 | 0.36 |
| 8 | switch | which location (of 9 options) | Day2_accDay1 | 0.30 | 0.46 |

### 3.1. More PFC activity for schema-congruent compared to schema-incongruent trials

First, we compared control trials (i.e., trials in which the event was updated with only schema-congruent information) and switch trials (i.e., trials in which the event was updated with schema-incongruent information). This revealed higher PFC activity for control trials compared to switch trials (when using aggregated data that is averaged across all 8 channels, see Figure 3). A linear mixed-effects model was conducted to examine how PFC activity varied during the encoding of control versus switch trials across the two days, which showed a significant interaction between *Trial Type* (control vs switch) and *Day* (day 1 vs day 2) (b = -5.00, SE = 0.98, t = -5.12, *p < .*001), which indicates that the increase in PFC activity from day 1 to day 2 was significantly smaller for switch trials (i.e., updating with schema-incongruent information) compared to control trials (i.e., updating with schema-congruent information). There was also a significant main effect of *Day* (b = 8.61, SE = 0.69, t = 12.46, *p < .*001), indicating greater PFC activity on day 2 compared to day 1, but no significant main effect of *Trial Type* (b = 0.46, SE = 0.69, t = 0.67, p = .51).

**Figure 3:**
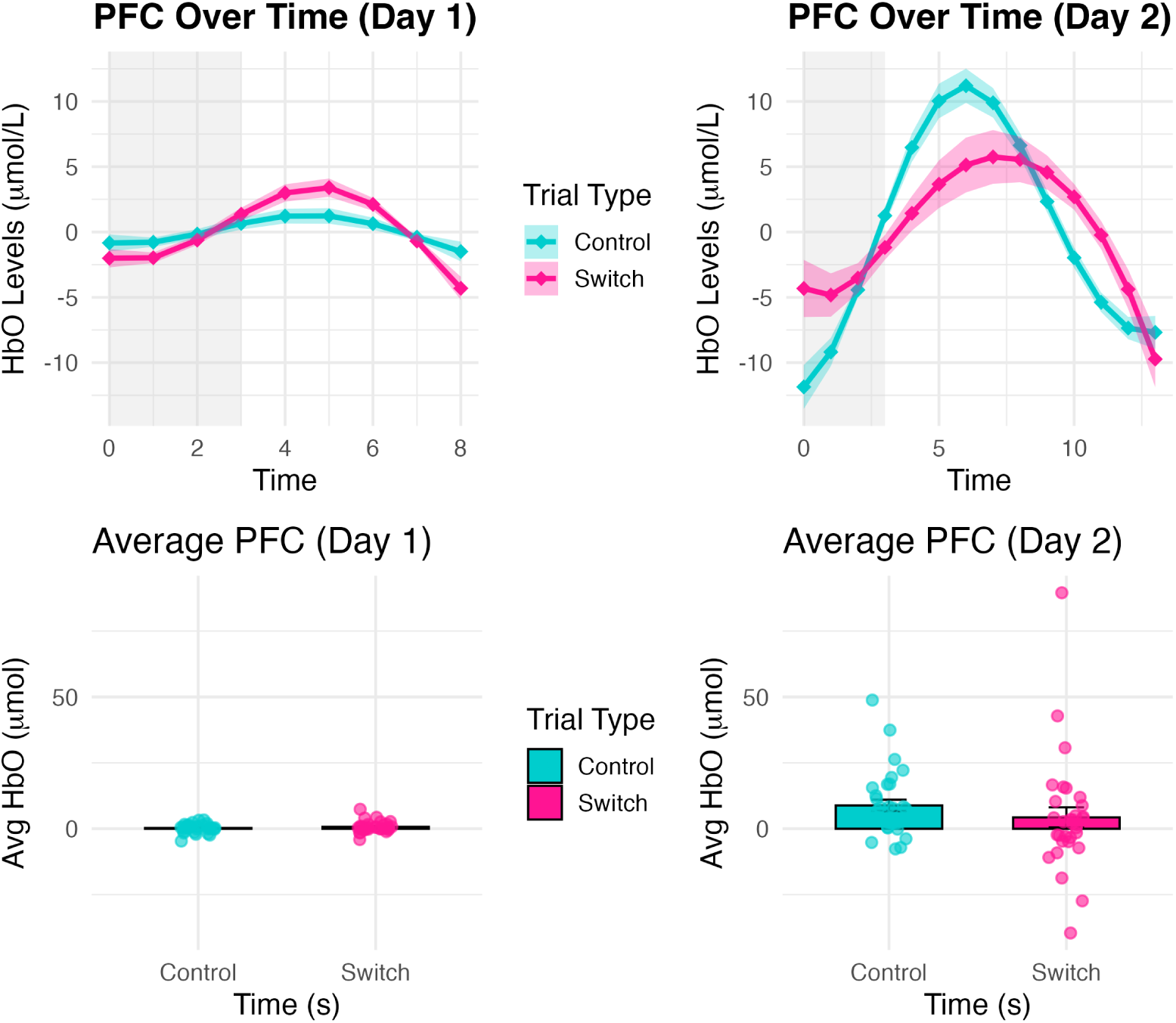
HbO levels (mean ± S.E.M.) from the PFC for Day 1 and Day 2 aggregated data (i.e., averaged across all 8 channels) in schema-congruent (control, blue) and schema-incongruent (switch, pink) conditions. Top left: Baseline Day 1 PFC activity for trials later classified as schema-congruent (blue) or schema-incongruent (pink). Top right: Day 2 PFC activity for schema-congruent and schema-incongruent trials. The shaded grey area marks the time window excluded from analysis due to the delayed onset of the hemodynamic response (HRF) underlying fNIRS signals. Bottom left: Mean HbO level on baseline Day 1 across time-points 4–8 s post-onset, corresponding to the expected peak of the BOLD response. Group mean ± S.E.M. is shown, with individual participant data overlaid. Bottom right: Mean HbO level on Day 2 across time-points 4–8 s post-onset, following the same conventions.

Next, we analyzed which sub-regions within the PFC were driving this effect of more activity during schema-congruent updating versus schema-incongruent updating (see Appendix Figure A.1). To get some indication for the differential contribution of PFC sub-regions, we labeled each of the 8 channels with its most likely PFC sub-region (labeling shown in Table 1). Channels most likely mapping onto the medial PFC (i.e., channels 1 and 5) showed this *Trial Type* * *Day* interaction (b = -8.09, SE = 2.17, t = -3.73, *p < .*001). Channels mapping most likely onto the Inferior Frontal Gyrus (i.e., channels 2 and 6) did not show a *Trial Type* * *Day* interaction (b = - 2.74, SE = 2.32, t = -1.18, p = 0.24). However, as shown in Appendix Figure A.1, channels 2 and 6 revealed a difference between the left (channel 2) and right (channel 6) Inferior Frontal Gyrus, with stronger PFC activity during control trials in the left and stronger PFC activity during switch trials in the right Inferior Frontal Gyrus. The remaining 4 channels (i.e., channels 3, 4, 7 and 8) all map most likely onto dorsolateral PFC, and all showed the same direction of stronger PFC activity in control compared to switch trials (b = -4.59, SE = 1.12, t = -4.11, *p < .*001).

In summary, medial and dorsolateral PFC regions as well as left inferior frontal gyrus all showed an increase in PFC activity for control trials (i.e., during updating with schema-congruent information) compared to switch trials (i.e., during updating with schema-incongruent information).

Day 1 data comparing control to switch trials is treated as a baseline for these analyses, since the difference between control and switch trials is only established on day 2, once the events become updated with additional information. Plotting the aggregated PFC data (i.e., averaging across all 8 channels) as well as the channel-wise data for day 1 confirmed that there is no meaningful difference yet on day 1 between later schema-congruent (control) versus later schema-incongruent (switch) trials (see Figure 3, left side and Appendix Figure A.2), as also evidenced by a linear mixed effects model on only day 1 data, comparing later-control to later-switch trials, that does not show a significant main effect of *Trial Type*: (b = 0.46, SE = 0.38, t = 1.20, p = 0.23) on baseline day 1. The same linear mixed effects model on only day 2 data, comparing control to switch trials, does indeed show a significant main effect of *Trial Type*: (b = -4.54, SE = 0.83, t = -5.46, *p < .*001).

### 3.2. Differences in PFC activity depending on whether or not participants re-integrated events with another schema context due to conflicting information

Then, we investigated differences between control trials (i.e., updating events with additional details while keeping the schema association the same) referred to as *schema-consistent* trials, successful switch trials (i.e., updating events with additional details while the participant successfully switched the schema association to the new, day 2 information) referred to as *schema-updating* trials, and unsuccessful switch trials (i.e., updating events with additional details while the participant still reported the original, day 1 schema association on day 2) referred to as *schema-violating* trials. Schema-consistent trials have the highest mean HbO levels (see Figure 4 and Appendix Figure A.4) in the time-period corresponding to the first 5 seconds of the trial, after excluding the beginning of the trial timeline due to the expected delayed onset of the HRF (see Methods), i.e. seconds 4 to 8. Schema-violating and schema-updating trials both showed a somewhat delayed peak (i.e., corresponding more to the last 5 seconds of the 14 second trials) relative to schema-consistent trials. Interestingly, while schema-violating trials seem to have a somewhat later peak of PFC activity that is somewhat lower in amplitude, schema-updating trials seem to first show a *decrease* in PFC activity after subsequently showing an increase in PFC activity towards the end of the trial (see Figure 4).

**Figure 4:**
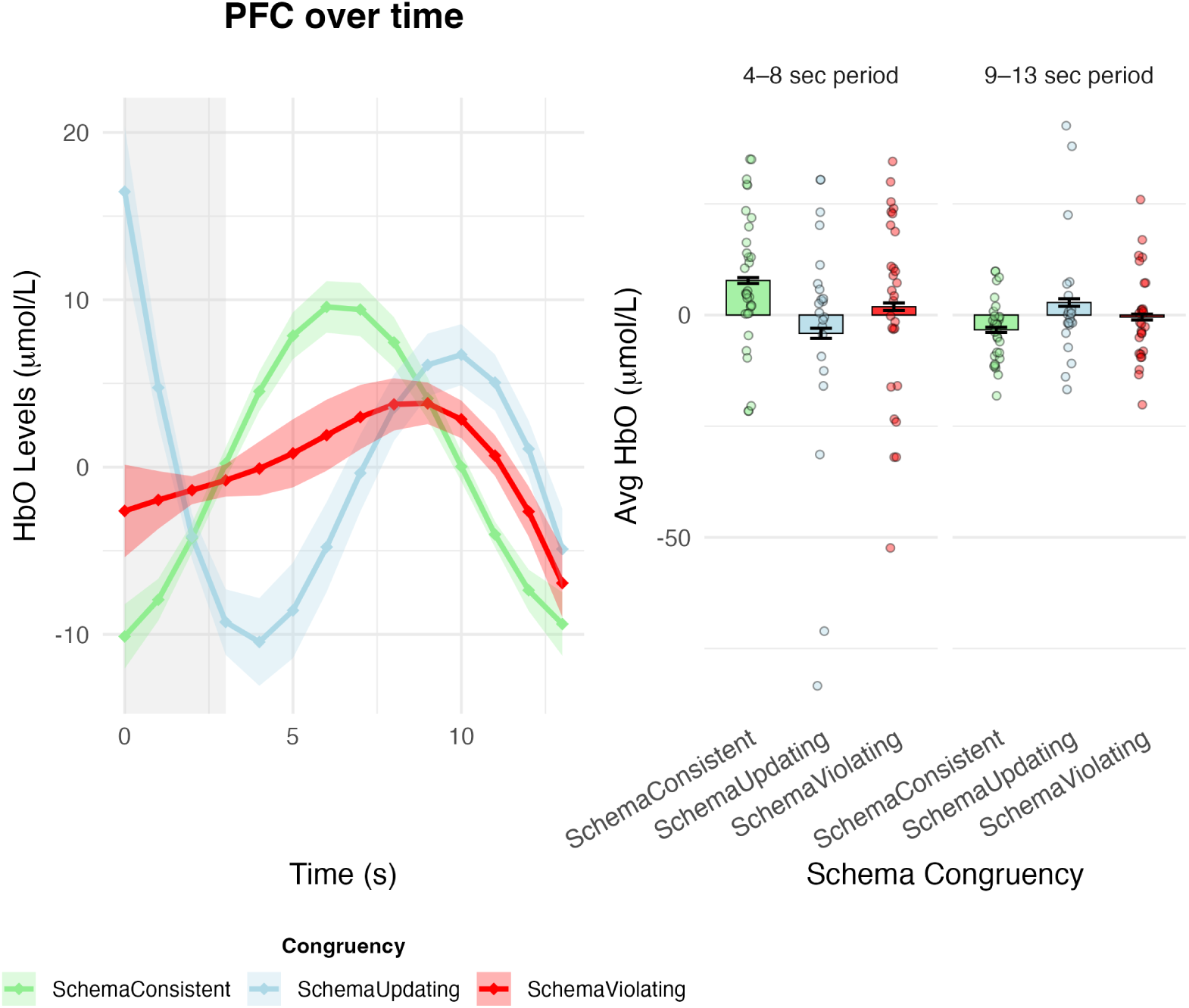
Aggregated HbO levels (mean ± S.E.M.) in PFC (i.e., averaged across all 8 channels) across schema consistent (green), schema violating (red) and schema updating (blue) conditions on day 2. Left: Entire timeline of the event (the shaded grey area marks the time window excluded from analysis due to the delayed onset of the HRF underlying fNIRS signals). Right: Mean ± S.E.M. overlaid with individual data points, averaged separately over seconds 4–8 and 9–13 post-stimulus onset.

Linear mixed-effects models were conducted to examine how congruency of the information (*Schema Consistent*, *Schema Updating*, *Schema Violating*) used for updating the events influenced HbO activity in the PFC in the period of the expected peak of the BOLD response (i.e., 4-8 seconds postonset). Schema-updating trials (b = -10.49, SE = 1.32, t = -7.97, *p < .*001) and schema-violating trials (b = -6.09, SE = 1.20, t = -5.09, *p < .*001) both showed lower PFC activity compared to schema-consistent trials. To investigate directly the difference between schema-updating and schema-violating trials, we followed up with another linear mixed-effects model that only included data from schema-updating and schema-violating trials, which showed a marginally significant difference (b = 3.03, se = 1.58, t = 1.92, p = 0.055). We again investigated which PFC sub-regions (mPFC, dlPFC and IFG) contributed most to this effect by analyzing channels mapping most likely onto the mPFC (i.e., channels 1 and 5), channels mapping most likely onto the IFG (i.e., channels 2 and 6) and channels mapping most likely onto the dlPFC (channels 3, 4, 7, and 8) separately. All 3 ROIs (likely mPFC, IFG and dlPFC) showed stronger PFC activity for both schema-updating and schema-violating trials in that first time-period (i.e., 4-8 seconds post-onset). In the likely-mPFC, schema-updating trials (b = -9.78, SE = 2.79, t = -3.50, *p < .*001) and schema-violating trials (b = -4.99, SE = 2.54, t = -1.96, p = 0.05) both showed lower PFC activity compared to schema-consistent trials. Also in the likely-IFG, schema-updating trials (b = -12.21, SE = 3.25, t = -3.76, *p < .*001) and schema-violating trials (b = -7.93, SE = 2.96, t = -2.68, p = 0.008) showed lower PFC activity compared to schema-consistent trials. Also in the likely-dlPFC, schema-updating trials (b = -10.29, SE = 1.44, t = -7.13, *p < .*001) and schema-violating trials (b = -5.61, SE = 1.31, t = -4.28, *p < .*001) both showed lower PFC activity compared to schema-consistent trials. The difference between schema-violating and schema-updating trials directly is most prominent in channels mapping onto the dorsolateral PFC (b = 3.87, SE = 1.62, t = 2.38, p = 0.017).

As can be seen in Figure 4, schema-updating and schema-violating trials seem to have a peak in PFC activity somewhat later in time in the trial compared to the schema-consistent trials. To investigate this, we conducted the same linear mixed-effects models but now on the next 5 seconds of the trial (i.e., second 9 to 13). The results showed that schema-updating (b = 5.84, SE = 1.11, t = 5.25, *p < .*001) and schema-violating (b = 2.83, SE = 1.01, t = 2.79, p = 0.005) trials both have higher PFC activity in the last 5 seconds of the trial compared to schema-consistent trials. A linear mixed-effects model that only included data from both schema-incongruent conditions showed a marginally significant difference with schema-updating having higher PFC activity then schema-violating (b = - 2.64, se = 1.36, t = -1.95, p = 0.052). When investigating PFC differences in these last 5 seconds of a trial for each of the 3 ROIs separately, we discovered that all three regions – likely-mPFC (b = 6.11, SE = 2.51, t = 2.43, p = 0.015), likely-dlPFC (b = 5.66, SE = 1.24, t = 4.56, *p < .*001), likelyIFG (b = 6.44, SE = 2.64, t = 2.44, p = 0.015) – showed higher PFC activity for schemaupdating then for schema-consistent trials in these last 5 seconds, but only the dlPFC also showed higher PFC activity for schema-violating compared to schema-consistent trials (b = 2.68, SE = 1.14, t = 2.36, p = 0.018).

## 4. Discussion

How is the PFC involved when new information comes to light that changes the schema-affiliation of earlier encoded events? Here, we ran a two-day fNIRS study with two goals in mind. Our first goal was to replicate what was found using fMRI (Bein and Niv, 2025; Masís-Obando et al., 2022; Baldassano et al., 2018; Gilboa and Marlatte, 2017; Brod et al., 2015; van Kesteren et al., 2012) regarding PFC involvement when updating with schema-congruent information with low-density and portable fNIRS equipment. Beyond mere replication, we wanted to find out whether fNIRS can be used in ecologically-valid and naturalistic memory paradigms. Our findings replicate previous fMRI findings. Schema-congruent information showed higher PFC activity than schema-incongruent information, but now with fNIRS. Our second goal was better understand PFC involvement in revising schemas. Our findings indicate that separating an event from its original schema and successfully re-integrating it into a new schema was characterized by an initial decrease in PFC activity early in the trial, followed by a later increase. This is notably different from trials where re-integration with a new schema failed and in which fNIRS recorded PFC activity had no initial decrease in PFC activity, and a lower-amplitude and later onset of PFC increase.

### 4.1. Updating with schema-congruent information

In general, we observed an increased PFC activity for schema congruent compared to schema incongruent events. This is in line with previous fMRI work on schema-congruency being largely dependent on (m)PFC (Bein and Niv, 2025; Audrain and McAndrews, 2022; Masís-Obando et al., 2022; Gilboa and Marlatte, 2017; Van Kesteren et al., 2013). To the best of our knowledge, there is no prior research on schema learning using fNIRS, but prior fNIRS studies do show involvement of PFC in associative learning (Jahani et al., 2017; Schaeffer et al., 2014), which is consistent with our findings given that schema learning builds on associative learning. This effect was most prominent in medial and dorsolateral PFC –and not in the IFG– which is in line with an abundance of fMRI research that altogether shows that medial PFC is involved when someone is confronted with schema-congruent information (Reagh and Ranganath, 2023; Masís-Obando et al., 2022; Raykov et al., 2021; van Kesteren et al., 2020; Baldassano et al., 2018; Preston and Eichenbaum, 2013; van Kesteren et al., 2010; Tse et al., 2007). Based on work from Brod et al. (2015), one would expect that channels mapping mostly onto the dorsolateral PFC would show an increase for updating with schema-incongruent information, due to the demands of cognitive control in these situations, with the assumption that the dorsolateral PFC is involved in conflict detection and subsequent resolution (Yang et al., 2024a; Oehrn et al., 2014; Wittfoth et al., 2009; Mansouri et al., 2009, 2007). However, our results show a different pattern. The dorsolateral PFC and the medial PFC both had increased activity for schema-congruent updating, especially in the left hemisphere.

There are several speculative reasons as to why this might be the case. Dorsolateral PFC is often suggested to be involved in post-retrieval monitoring and evaluation, especially in situations in which retrieval of episodic information involves high ambiguity and/or high demands on verification (Hayama and Rugg, 2009). Thus, for tasks in which one must evaluate features of earlier memory content to make some kind of decision. In schema-congruent updating trials, participants may closely monitor whether a given detail reflects a true memory from the original Day 1 description, a schema-driven inference, or a genuinely new detail introduced on Day 2. Given the conflict between the new details and their prior knowledge in schema-incongruent trials, participants might be less engaged with deciding how the new details fit with the prior event description compared to schema-congruent trials. Due to the high similarity and potential overlap between schema-congruent new details and existing memory traces, schema-congruent trials might rely more heavily on source evaluation –a process that engages metacognitive monitoring (Vaccaro and Fleming, 2018; Fleming and Dolan, 2012). This, in turn, is thought to depend on the dorsolateral PFC, which supports the controlled evaluation of memory sources (Saccenti et al., 2024). Additionally, a recent meta-analysis revealed dorsolateral PFC to be consistently involved in temporal context retrieval, for which one would need to organize a sequence of related events in order to determine how these relate to each other in terms of time (Torres-Morales and Cansino, 2024). Interjecting these added details on day 2 of our experiment in the original sequence of information of an event might in particular have been relevant for schema-congruent updating. Therefore, updating with schema-congruent information might still (partly) rely on the dorsolateral PFC as well as the medial PFC in our study.

### 4.2. Dissociating events from one schema and subsequently integrating it with another schema

Both schema-incongruent conditions, schema-updating and schema-violating, leads to a conflict between the events shown on day 1 and the new details added on day 2. Conflict detection is known to involve PFC (Yang et al., 2024a; Li et al., 2017; Zmigrod et al., 2016; Oehrn et al., 2014; Sun et al., 2013). However, updating with new details in both schema-incongruent conditions seem to lead to a peak in PFC activity *later in time* compared to schema-consistent control trials. One possible interpretation of this temporal pattern is that the conflict and resulting prediction error gradually build over the course of the trial, as the mismatch between incoming information and existing memory representations becomes more evident. In schema-incongruent trials—both schema-updating and schema-violating—participants are likely primarily engaged in detecting this conflict between the new information and their prior knowledge, which dominates early processing stages (Richter et al., 2016; van Kesteren et al., 2012). Updating of the memories may only occur once the conflict is brought to conscious awareness, highlighting the role of metacognitive monitoring that enable individuals to detect inconsistencies and adaptively revise their memory representations (Boldt and Gilbert, 2022; Chua et al., 2014). The temporal accumulation of conflict may delay the engagement of higher-order cognitive control mechanisms, resulting in the later peak in PFC activity observed in these trials. Later PFC recruitment likely reflects the time required to override conflicting schema-driven expectations and to re-evaluate the event in light of the unexpected information. This process is fundamentally different from schema-consistent control trials, where updating is fully schema-driven, which is known to be quicker and easier (Coutanche and Thompson-Schill, 2014; Tse et al., 2007). An interesting finding in our study is that those trials for which updating with schema-incongruent details subsequently led to successful re-integration of the event with another schema showed an initial dip in PFC engagement, prior to the peak later in time. We speculate that the initial dip in PFC activation may not simply reflect a lack of engagement, but rather a controlled disengagement from the existing schema-based representation. In order to successfully update an existing Day 1 memory with incongruent Day 2 information, participants might first need to suppress or inhibit the influence of prior knowledge, thereby creating cognitive space for restructuring the event representation. This transient reduction in PFC activity may thus be a functional mechanism that enables re-encoding of the new information. This would be in line with theories about memory updating that indicate that prediction errors – which would be stronger for schema-incongruent trials in our experimental design – would serve as cues to weaken a memory representation in order to update memories (Wahlheim and Zacks, 2025). Empirical evidence in this direction can be found in studies using directed forgetting paradigms that reveal that instructing participants to forget an earlier memory can lead to weakening of earlier memories and strengthening of new memories (Chiu et al., 2021; Manning et al., 2016).

In this interpretation, the absence of an initial dip, such as in schema-violating trials where no successful updating occurs, might indicate that the prior schema remains continuously active. If the PFC immediately ramps up to resolve conflict without first quieting the initial schema association, it may reinforce competing representations, leading to interference between Day 1 and Day 2 memories (Gilboa, 2019; Runquist, 1975). Therefore, we speculate that the initial dip may play a critical regulatory role in enabling effective memory updating by allowing for adaptive schema disengagement at first, followed by strategic re-encoding processes supported by later PFC activation.

### 4.3. Future research directions

Future research could further validate and expand on these findings. First, follow-up studies using high-density fNIRS systems could enhance spatial resolution, enabling the investigation of schema updating processes in a more fine-grained manner. Second, complementary fMRI studies are needed to explore the interplay between the PFC and medial temporal lobe (MTL) regions (Gilboa and Marlatte, 2017; Van Kesteren et al., 2013) –particularly the hippocampus– during schema revision and re-integration, to better characterize the dynamics between PFC and MTL underlying dissociation and re-integration of events into schemas. Third, although multivariate pattern analysis (MVPA) in the PFC presents known challenges (Bhandari et al., 2018), future attempts to decode schema-related neural patterns could reveal finer-grained distinctions between successful and failed schema revision, either with fMRI or with high-density fNIRS. Finally, computational modeling approaches could be used to shed light onto whether our interpretations of the observed pattern in the schema-updating condition of early PFC deactivation followed by a later peak indeed reflects a process of initial schema suppression followed by successful re-integration into an updated knowledge structure.

## 5. Conclusion

The fNIRS data revealed stronger PFC activation when events were updated with schema-congruent rather than schema-incongruent information. This finding aligns with previous fMRI studies and supports the use of fNIRS as a dependable tool for memory research, particularly in ecologically valid and naturalistic settings. For events involving schema-incongruent updates, we observed an initial drop in PFC activity early in the trial, followed by a later increase, suggesting a two-step process of detaching the event from its original schema and re-integrating it into a new one. This temporal pattern differed notably from trials where re-integration failed, which did show a similar delayed PFC increase, albeit lower in amplitude, and did not show an initial decrease in PFC activity at all. Overall, these findings offer a more nuanced understanding of how the brain flexibly updates memories when faced with conflicting information. They highlight the dynamic role of the PFC in detecting discrepancies and restructuring memory accordingly. This has important implications for understanding false memories: when new details that are incongruent with earlier knowledge are presented, the brain may either successfully update the earlier memory or not. Gaining deeper insight into the mechanisms that determine whether updating occurs can help explain why certain situations are more prone to the incorporation of misleading or inaccurate details. Our results suggest that both the timing and the strength of PFC engagement may determine whether conflicting information is incorporated or rejected. These insights contribute to a deeper understanding of the neural mechanisms underlying memory distortion and the challenges of correcting misinformation.

## Author CRediT statement

**Petra Biro**: Conceptualization, Software, Data Curation, Investigation, Validation, Formal analysis, Visualization, Writing - Original Draft. **Silvy H.P. Collin**: Conceptualization, Formal analysis, Visualization, Supervision, Project administration, Writing - Review and Editing.

## Acknowledgements

We thank members of the DAF Technology lab at which we ran data acquisition of this project. In particular, we thank Hans van den Dool for help regarding IT and fNIRS hardware/software set-up in the fNIRS lab.

## Appendix

**Figure A.1:**
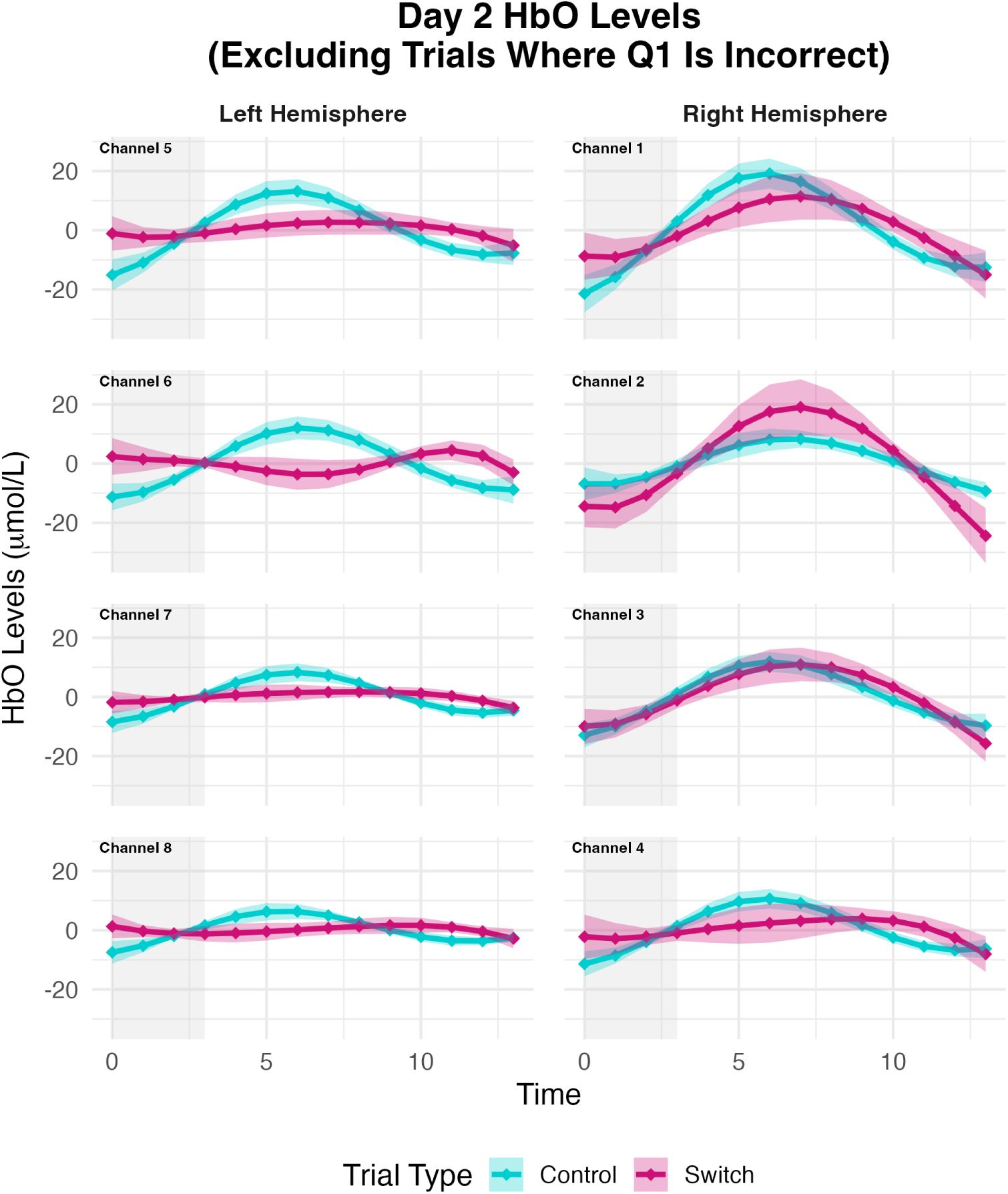
Channel-wise HbO levels for schema congruent (control) and schema incongruent (switch) trials on day 2. Only trials included were people answered the old/new question correctly (i.e., remembered trials). Channels 1 and 5 most likely map onto the mPFC, channels 2 and 6 most likely onto the IFG, and channels 3, 4, 7 and 8 most likely onto the dlPFC.

**Figure A.2:**
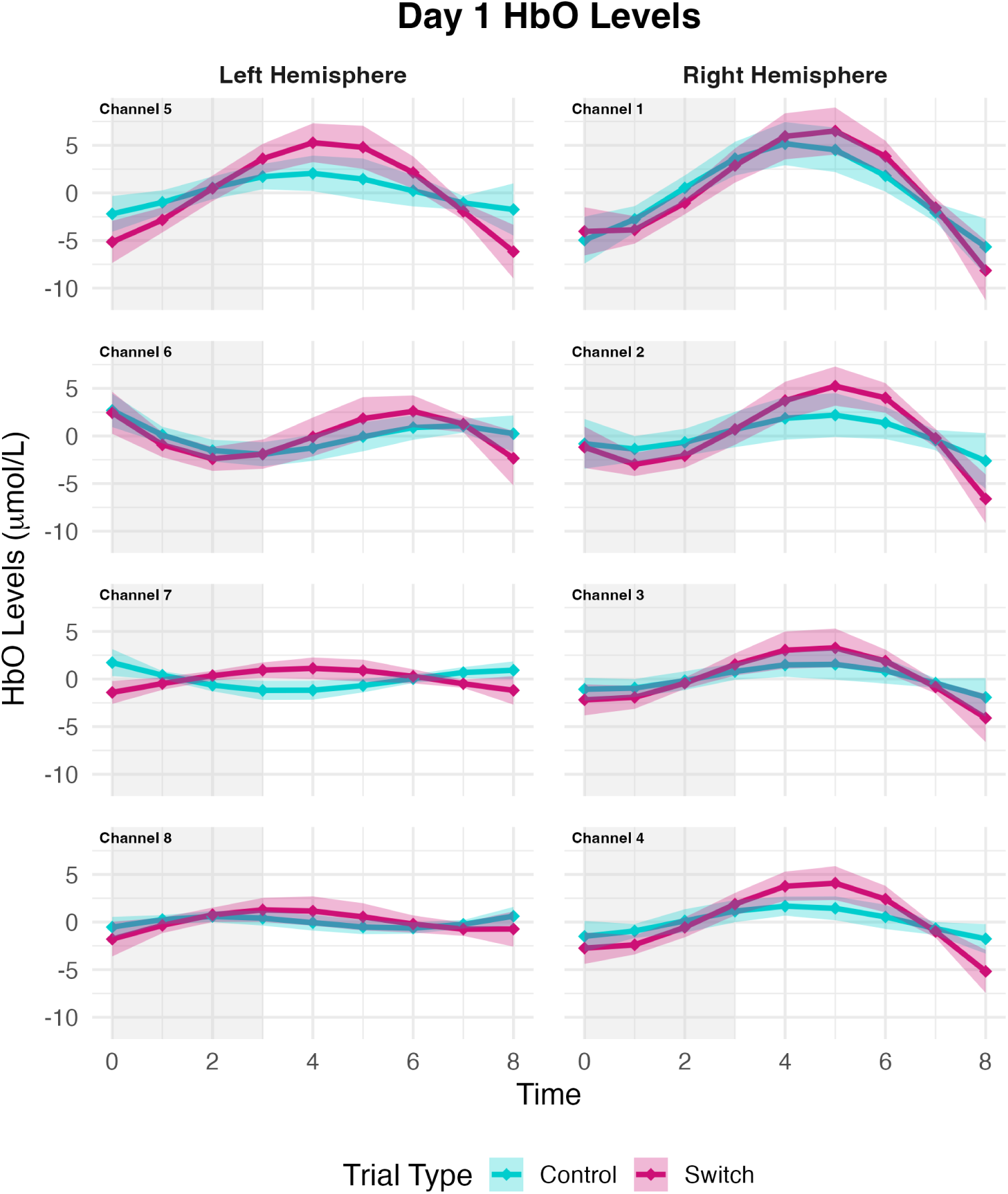
Channel-wise HbO levels for later schema congruent (control) and later schema incongruent (switch) trials on day 1. Channels 1 and 5 most likely map onto the mPFC, channels 2 and 6 most likely onto the IFG, and channels 3, 4, 7 and 8 most likely onto the dlPFC.

**Figure A.3:**
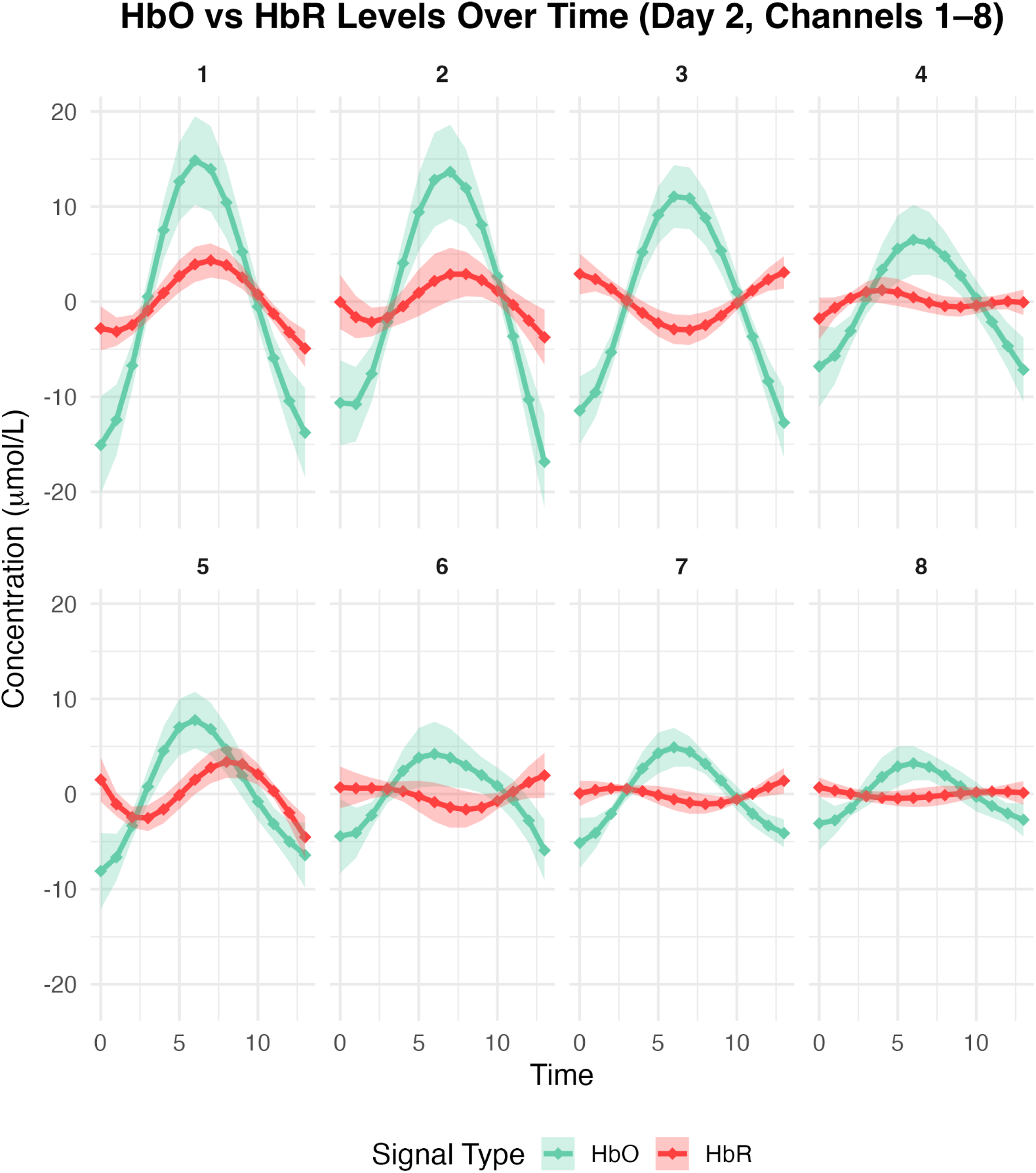
Average HbO versus HbR per channel, for day 2.

**Figure A.4:**
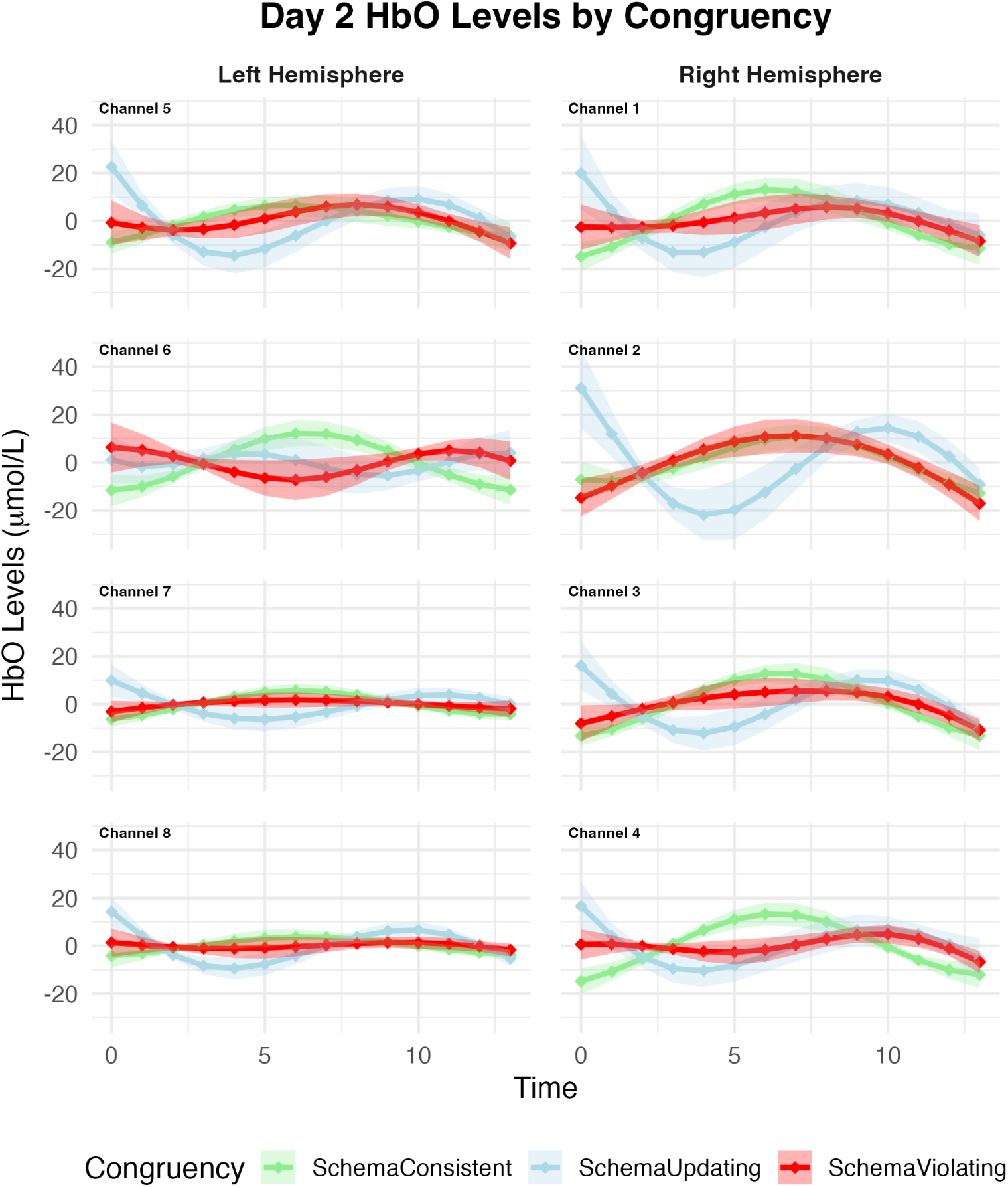
Channel-wise HbO levels (mean ± S.E.M.) across schema consistent (green), schema violating (red) and schema updating (blue) conditions on day 2. Channels 1 and 5 most likely map onto the mPFC, channels 2 and 6 most likely onto the IFG, and channels 3, 4, 7 and 8 most likely onto the dlPFC.

## Notes

### Competing Interest Statement

The authors have declared no competing interest.

